# Antibody Profiles in Pediatric Autoimmune Neuropsychiatric Disorders Associated with Streptococcal Infections

**DOI:** 10.64898/2026.05.11.724168

**Authors:** Thomas J. Esparza, Nicole F. Lee, Margaret Pekar, Pavel P. Khil, Christopher M. Bartley

## Abstract

Pediatric Autoimmune Neuropsychiatric Disorders Associated with Streptococcal Infections (PANDAS) is characterized by prepubertal abrupt onset of obsessive-compulsive disorder (OCD). The sine qua non is group A streptococcus (GAS) infection, which is hypothesized to elicit an IgG-class anti-GAS antibody response that cross-reacts with antigens in the basal ganglia. However, the association between GAS antibody (GAS-IgG) levels and PANDAS has been inconsistent, and qualitative differences in GAS-IgG profiles have not been carefully evaluated in well-phenotyped cohorts. Moreover, independent studies have yet to converge on anti-neural autoantibodies that are specific to PANDAS.

Here, we used phage display immunoprecipitation sequencing (PhIP-Seq) to perform ultra-deep anti-pathogen antibody repertoire profiling of serum from definitive pediatric PANDAS patients (N = 34) collected as part of a prior double-blind, placebo-controlled clinical trial of intravenous immunoglobulin (IVIg). PANDAS cases were compared to pediatric controls without a history of neuropsychiatric illness (N = 31). To assess for objective evidence of neuroglial injury, serum neurofilament light (NfL) and glial fibrillary acidic protein (GFAP) levels were compared to healthy pediatric controls. Within PANDAS, NfL and GFAP levels were compared between pre- and post-treatment sera. To evaluate for central autoantibodies, a subset of baseline cerebrospinal fluid (CSF) samples (N = 25) was profiled by full-length human protein microarray.

Though GAS reactivity by PhIP-Seq was well correlated with clinical anti-DNaseB and anti-streptolysin O titers, there were no quantitative or qualitative differences in GAS-IgG profiles between PANDAS and controls. Furthermore, NfL and GFAP levels did not differ between cases and controls. Within PANDAS, changes in NfL or GFAP levels at six weeks did not differ between placebo and IVIg groups. However, CSF autoantibody profiling by protein microarray revealed infrequent but notable candidate autoantibodies. In one patient, we identified autoantibodies against Argonaute family proteins (AGO-IgG), a marker of autoimmune sensory neuropathy. Longitudinal measurement of AGO-IgG in sera revealed that titers were unchanged after placebo, but decreased after IVIg, coinciding with symptomatic improvement, including a decrease in that patient’s CY-BOCS score.

Overall, these results do not support an etiologic role for GAS-IgG in PANDAS. However, some individuals diagnosed with PANDAS may harbor anti-neural autoantibodies.

## Introduction

Pediatric Autoimmune Neuropsychiatric Disorders Associated with Streptococcal Infections (PANDAS) is primarily characterized by prepubertal acute-onset obsessive-compulsive symptoms (OCS) associated with *Streptococcus pyogenes* (group A *Streptococcus* [GAS]), a globally endemic bacterium that causes acute upper respiratory tract and skin infections. GAS infection can be complicated by immune-mediated rheumatic fever, rheumatic heart disease, post-streptococcal glomerulonephritis, and Sydenham chorea (SC). However, a meaningful proportion of individuals with SC also exhibit neuropsychiatric features, including OCS (*1, 2*), which have also been reported in rheumatic heart disease without SC (*3*).

PANDAS is hypothesized to arise from molecular mimicry, whereby anti-GAS antibodies (GAS-IgG) cross-react with an unidentified autoantigen in the basal ganglia (*4*), similar to the molecular mimicry that underlies valvular disease in rheumatic heart disease (*5*). In PANDAS, anti-lysoganglioside (*6, 7*), tubulin (*8*), and other antibodies (*9*) have been reported as the antigenic targets. However, one prospective longitudinal study failed to identify a correlation between GAS-associated symptom exacerbation and lysoganglioside reactivity (*10*). Subsequent studies failed to find a strong association between GAS and PANDAS (*11, 12*).

As a result, patients who met criteria for PANDAS but whose symptoms were not clearly associated with GAS posed a diagnostic conundrum. PANDAS was therefore reframed as a subsyndrome of pediatric acute-onset neuropsychiatric syndrome (PANS) (*13*), which allowed for non-infectious and infectious triggers, including *Mycoplasma pneumoniae (M. pneumoniae), Chlamydia pneumoniae (C. pneumoniae*), Epstein-Barr virus (EBV), *Borrelia burgdorferi* (Lyme disease), and herpes simplex virus 1 (HSV-1) (*14*). PANS further dispensed with the requirements of prepubertal onset while requiring at least two additional neuropsychiatric symptoms: anxiety, emotional lability or depression, irritability/aggression/oppositional behavior, behavioral regression, deterioration in school performance, sensorimotor abnormalities, or somatic signs (sleep changes, enuresis, or urinary urgency). Nonetheless, the diagnosis of PANDAS remains in clinical practice, with reports that GAS titers distinguish between PANDAS and PANS (*15*).

Therefore, whether PANDAS is defined by a unique GAS-IgG response remains unknown and has implications for the validity of PANDAS as a singular disease entity. Notably, up to 75% of individuals diagnosed with PANDAS fail to meet full PANDAS criteria (*16*). Therefore, we profiled archived samples from a prior double-blind placebo-controlled clinical trial of intravenous immunoglobulin (IVIg), wherein 36 of 1,183 screened subjects ultimately met strict criteria for PANDAS and were enrolled. GAS-IgG profiles were evaluated by a phage display immunoprecipitation sequencing (PhIP-Seq) library encoding ∼753,000 peptides derived from parasites, fungi, viruses, and bacteria, including GAS. Sera were further evaluated for elevated neural injury markers by Quanterix Simoa and CSF for anti-neural autoantibodies by protein microarray.

## Results

### Cohort and Study Design Characteristics

Detailed demographic data are available in the original study (*17*). Briefly, we summarize the age, sex, and demographics of the enrolled PANDAS cohort. A total of 48 participants completed in-person assessments at NIMH, including 27 males and 21 females, ranging in age from 4 to 12 years. Of these, 36 participants were enrolled in the IVIg trial. Of these participants, 24 were male, and 12 were female, with a mean age of 9.26 ± 2.32 years (range, 4 - 12). The mean baseline CY-BOCS score was 27.64 ± 4.59, indicating moderately-severe OCS. Subjects underwent serum and CSF sampling at baseline prior to administration of placebo (N = 18) or IVIg (N = 17). One randomized participant (IVIg) withdrew from the study prior to study drug infusion. Subjects underwent repeat serum and CSF sampling 6 weeks after treatment. After unblinding, the study converted to open-label optional initiation or continuation of IVIg and repeat serum sampling at 12 and 24 weeks.

### PhIP-Seq profiling of IgG reactivities in PANDAS

To comprehensively characterize IgG class immunoreactivity towards microbial antigens in PANDAS patients, we performed PhIP-Seq using the PanSeq exposure library component (*18*). The PanSeq exposure libraries consist of a broad representation of both bacterial and viral proteins and other pathogen peptides across ∼753,000 total peptides.

Following immunoprecipitation (IP) and sequencing, reads were aligned to the reference oligo library. Strongly reactive peptides were identified by comparing to mock IP, input library, and low reactivity sera. In total across cases and controls, 19,217 peptides were enriched in at least one sample by >4-fold relative to mock IP samples, input libraries, and low-reactive sera. Overall, between 600 and 4382 reactive peptides per patient were identified (median = 1,090, IQR: 878-1,310; **Figure 1A**).

**Figure 1.**
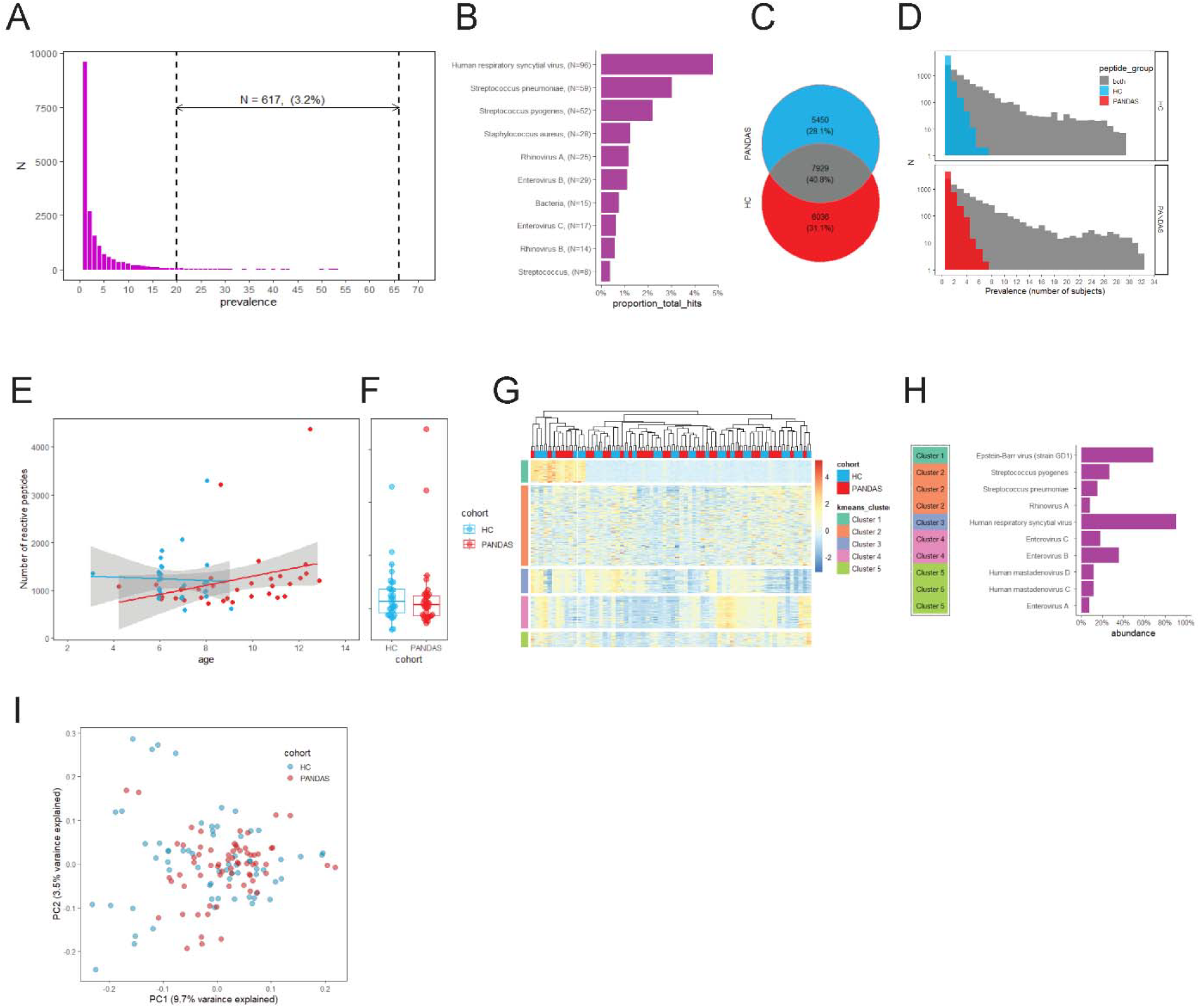
PhIP-Seq profiling of anti-microbial IgG response in HC and PANDAS cohorts. (A) Histogram of population prevalence of enriched peptides. Majority of peptides are detected in one or two subjects. (B) Taxonomic composition of most prevalent (N>20) PhIP-Seq reactivities. Top 10 taxonomic units among broadly reactive peptides are shown. (C) Summary of enriched peptide detection in HC and PANDAS. (D) Histogram of population prevalence of enriched peptides in each of the study cohorts. All peptides are separated in groups of peptides detected in both cohort or only in one. (E) Dependence on age on immune response diversity. Number of enriched peptides is plotted for each cohort along with linear regression fit. Grey area represents 95% CI area for regression coefficient. (F) Boxplots and points plot of number of enriched peptides separated by study cohort, (G) Clustered heatmap of peptide reactivities (top 300 peptides with largest variance). Peptide reactivity matrix was log2-transformed and re-centered row-by-row prior to plotting and analysis. Samples are annotated with sample cohort assignment and peptides with k-means cluster. Columns were ordered using hierarchical clustering prior to plotting. (H) Taxonomic composition of k-means clusters. Only taxonomic units with >10% of each cluster content are included. (I) PCA plot of study samples. Analysis is restricted to set of 5543 peptides detected in 3 or more samples.

Consistent with the high variability of individual immune responses and the distinct exposure history of study subjects, most of the enriched peptides were restricted to relatively few subjects. Overall, 49.5% (N = 9,503) of all peptides are enriched in a single subject, and 3.4% (N = 653) were detected in 20 or more subjects (**Figure 1A**). This highly concentrated pattern of reactivities is consistent with earlier studies where anti-viral response in adults was studied (*19*). Enriched peptides with the highest prevalence map to common pathogens, including respiratory syncytial virus (RSV) (N=92 peptides with prevalence above 20), *S. pneumoniae* (N=57), and *S. pyogenes* (N=50, **Figure 1B**). Yet, even the most highly prevalent reactivities account for less than 2%-5% of all enriched peptides (**Figure 1B**).

Strongly reactive peptides were distributed evenly between study cohorts. 13,379 peptides were detected in PANDAS, 13,651 in HC, whereas 7,813 (40.7%) of all enriched peptides were detected in both cohorts (**Figure 1C**). Importantly, those peptides detected in PANDAS and HC cohorts had a significantly higher mean prevalence than those detected in only one cohort (mean prevalence = 4.1 versus 1.2 in HC- and 1.3 in PANDAS-only peptides, P < 2.2e-16, one-way ANOVA, **Figure 1D**).

There was also no difference in the diversity of IgG reactivity profiles represented as the number of enriched peptides in PANDAS compared to HC (**Figure 1F**, P=0.84, t-test). Additionally, we did not detect a significant increase in diversity of immune response with age (**Figure 1E**). Unlike previous studies that reported a relationship between age and anti-viral antibody diversity (*20*), the age range of our subjects is very narrow, 3-13 years.

To characterize the pathogen response in our cohorts more deeply, we performed K-means clustering (N = 5) of reactivity profiles toward the 300 most variable peptides (**Figure 1G**). Clustering revealed several diverse patterns of reactivity. While for clusters 2-4 response patterns are largely similar, cluster 1 appears quite heterogeneous, suggesting that additional analysis may reveal finer structure. Taxonomic annotation (**Figure 1H**, species with >5% prevalence are shown) indicated that the primary determinants of clustering are common pathogens.

In agreement with the high similarity of response profiles within clusters 3 and 4, each is strongly dominated by a single species, 92% of peptides map to RSV in cluster 3 and 70% to EBV in cluster 4. Though less homogeneous, cluster 5 peptides predominantly (88%) mapped to *S. pyogenes*. Whereas cluster 2 represents Enterovirus B and C reactivity profiles, cluster 1 is substantially more diverse. The most frequent species in cluster 1, *S. pneumoniae*, accounted for only 14% of all peptides. As before, we did not observe an obvious difference in the distribution of study cohorts relative to pathogen response profiles (**Figure 1G**).

Finally, to evaluate the separation between HC and PANDAS subjects, we performed principal component analysis (PCA). For PCA, we used 54541 peptides enriched in 4 or more subjects (**Figure 1I**). Overall, there was no separation between PANDAS and healthy controls on the PCA plot, though the proportion of variance explained by PC1 and PC2 is 9.7% and 3.5%, respectively.

Taken together, although we detected diverse and abundant antimicrobial responses by PhIP-Seq, there were no obvious differences between HC and PANDAS groups.

### Association between PANDAS and GAS: an in-depth evaluation

First, we evaluated the distribution of enriched GAS peptides (defined as log_2_ fold change (LFC) >2, P.adj<0.01; where LFC = log_2_ fold change) in PANDAS and HC cohorts. In total, 204 GAS peptides were enriched in at least one study subject, with a mean of 42 peptides per subject (range 0 - 78, median = 41, **Figure 2A**). All subjects except for one HC individual had at least one enriched peptide (**Figure 2B**), and the proportion of reactive subjects for each peptide was roughly similar between HC and PANDAS. We then directly compared the frequency of reactivity to each peptide in HC and PANDAS cohorts (**Figure 2C**) and found that they were tightly correlated. Thus, taken together, there was no detectable difference in the broadly reactive immune response to GAS between PANDAS and HC.

**Figure 2.**
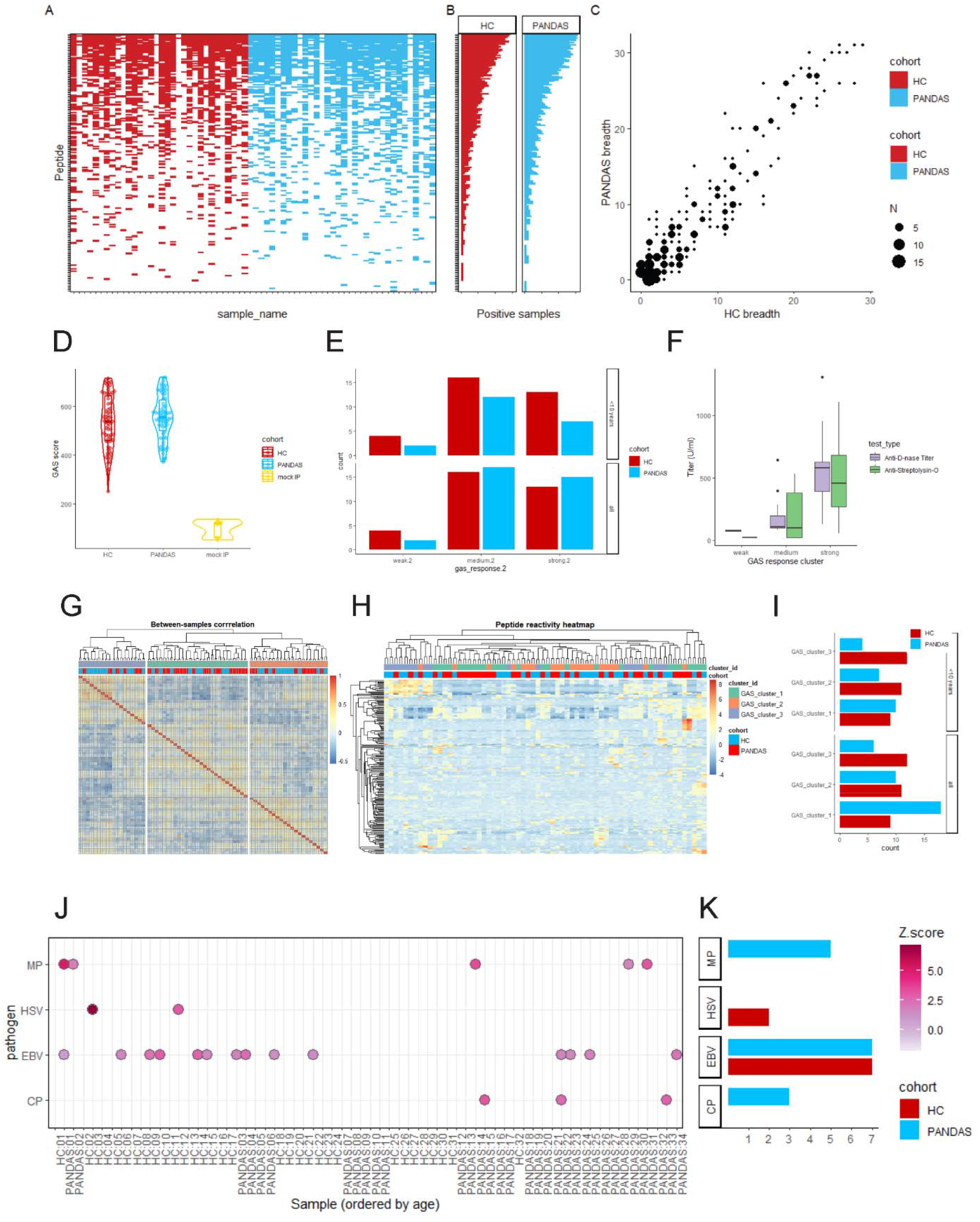
Association between pathogen reactivities and PANDAS. (A) Detection of individual enriched GAS peptides and (B) histogram of peptide prevalence across study cohort. Peptides are ordered by prevalence from top to bottom. (B) Comparison of GAS peptide prevalence in HC (x axis) and PANDAS (y axis). Number of peptides for each frequency combination is represented as circle size. Enriched peptide prevalence in PANDAS is tightly correlated to prevalence in HC. (D) Aggregated GAS score distribution in HC, PANDAS and mock IP samples. (E) Frequency of samples from total GAS reactivity class parsed by study cohort. Analysis was performed separately for all samples or subset of subjects < 10 years of age. (F) Anti-streptolysin and anti-DNase antibody titers determined by ELISA. All titers are parsed by GAS reactivity group. (G) Clustered heatmap of Pearson correlation coefficient for study samples. Samples are annotated with GAS response cluster assignment and study cohort. (H) Clustered heatmap of GAS peptide reactivities (top 200 most variable peptides). Both peptides (rows) and samples (columns) were clustered prior to plotting. Normalized peptide reactivities were log2-transformend and re-centered prior to plotting and analysis. Samples are annotated with cohort assignment and GAS response cluster. (I) Frequency of samples from each GAS response cluster parsed by study cohort. Analysis was performed separately for all samples or subset of subjects < 10 years of age. (J) Selected pathogen positivity as determined from PhIP-Seq data. Z-score was calculated relative to aggregate pathogen score reactivity across all samples. (K) Frequency of pathogen-positive samples parsed by study cohort. Analysis was performed separately for all samples or subset of subjects < 10 years of age.

Since the above analytic approach is restricted to strongly enriched peptides and can be less sensitive, we utilized an alternative strategy to evaluate exposure history and immune response to GAS. We identified all *S. pyogenes*-related peptides in our PanSeq exposure libraries and evaluated their reactivity. In addition to parsing taxonomy annotation, we performed a direct BLAST search using our reference proteome. In total, we detected 953 *S. pyogenes*-related peptides (**Supplementary Figure 1**, Supplementary Methods for details on peptide identification).

We then calculated the aggregate GAS reactivity score by summing the normalized reactivity values for all peptides. We found that the GAS reactivity score is significantly higher (mean+3xSD) than mock IP score for all study subjects (**Supplementary Fig 2**), both from HC and PANDAS cohorts (**Figure 2D**), but there is no difference in GAS score between HC and PANDAS (P=0.37, t-test).

Using this aggregate score, we subdivided all subjects into three groups based on GAS response—weak, intermediate, and strong—and asked whether higher or lower GAS reactivity scores are non-randomly distributed between study cohorts (**Figure 2E**). There is no difference in association between GAS response class cohort (chi-square test, X-squared = 1.70, P=0.43, all subjects). This lack of association was not due to a skew in age (chi-square test, X-squared = 0.84, P=0.66, age <10 years). Furthermore, GAS antibody titers have been reported to be relatively stable between ages 2 and 12 (*21*).

Next, we compared our PhIP-seq-derived GAS response score to historical DNase B and anti-streptolysin O (ASO) clinical antibody titers for these subjects (**Figure 2F**). We found that although there is categorical agreement, and subjects assigned to the high reactivity GAS cluster tend to have higher DNase and Streptolysin titers, there was still considerable variance (**Figure 2F, Supplementary Fig 3**).

Finally, we performed a deeper evaluation of the similarity between GAS reactivity profiles in our subjects. Using hierarchical clustering over the sample correlation matrix, we separated samples with similar reactivity profiles into three groups (**Figure 2G**). A detailed examination of the peptide reactivity heatmap indicates some level of heterogeneity in the immune response to GAS, perhaps driven by strain differences, and suggests that there could be further subdivision between samples (**Figure 2H**). Yet, when we similarly compared relative frequencies of GAS response clusters in study cohorts, we found no association (**Figure 2I**) (chi-square test, X-squared = 4.25, P=0.12, all subjects, X-squared = 1.64, P=0.44, <10 years).

Our focused analysis suggested that all or nearly all subjects have some measurable level of reactivity to GAS antigens.

### Association of other pathogens with PANDAS: *C*. *and M*. *Pneumoniae*

Because there was no apparent increase or difference in immune reactivity to GAS peptides in PANDAS. Since it has been reported that several other pathogens were associated either with PANDAS or PANS (*14, 15*), we asked whether IgG reactivity to these pathogen-encoded peptides is altered in PANDAS patients.

In total, we identified 241 *M. pneumoniae*-derived peptides, 221 *C. pneumoniae* peptides, 2,422 EBV peptides, and 2,368 HSV-1 peptides in the PanSeq exposure library. To calculate the reactivity score, we restricted the peptide set to those at least moderately reactive (max normalized reactivity > 2 cpk in our dataset; where cpK = read counts per 100,000) and then aggregated reactivity values as before. Using a cutoff of Q3 + 1.5 * IQR in both technical replicates, we identified pathogen-positive samples (**Figure 2J and Supplementary Figures 4-7**). While nearly all samples were GAS-IgG positive, reactivity to other pathogens was substantially lower. The highest positivity rate was observed for EBV at ∼20% (**Figure 2J, 2K**).

When we compared the pathogen positivity rate between PANDAS and HC cohorts, none of the rates were significantly different. Notably, the pathogen with the strongest imbalance, *M. pneumoniae* (N=5 PANDAS, N=0 HC positives, **Figure 2K**), nearly reached the significance threshold, yet it did not exceed it (Fisher’s Exact test, P=0.055). Thus, although there are some indications for the association of pathogen exposure to *M. pneumoniae* or perhaps *C. pneumoniae* with PANDAS, these warrant further investigation and replication.

### Neuroglial Injury Markers

PANDAS has been described in some studies as a form of encephalitis (*22, 23*). Though PANDAS lacks the classical radiographic features of encephalitis (*24*), encephalitis in the absence of radiographic changes has been well described (*25*). Therefore, we evaluated serum levels of neurofilament light (NfL) and glial fibrillary acidic protein (GFAP), sensitive biomarkers of neural injury that have been shown to discriminate between autoimmune encephalitis and primary psychiatric disorders (*26, 27*).

At the cohort level, there was no difference in baseline NfL or GFAP levels between PANDAS and controls (**Figure 3A**). Because NfL and GFAP levels change throughout adolescence, we evaluated levels relative to age-adjusted international reference ranges (*26, 28*). However, for both controls and PANDAS, nearly all levels fell within expected reference ranges (**Figures 3B and C)**. Next, we evaluated whether there was a change in NfL and GFAP levels between placebo- and IVIg-treated PANDAS subjects between baseline and 6 weeks. However, there was no effect of either treatment and mean changes did not differ between groups, indicating the absence of subclinical IVIg-responsive neural injury in PANDAS (**Figures 3D - F**).

**Figure 3.**
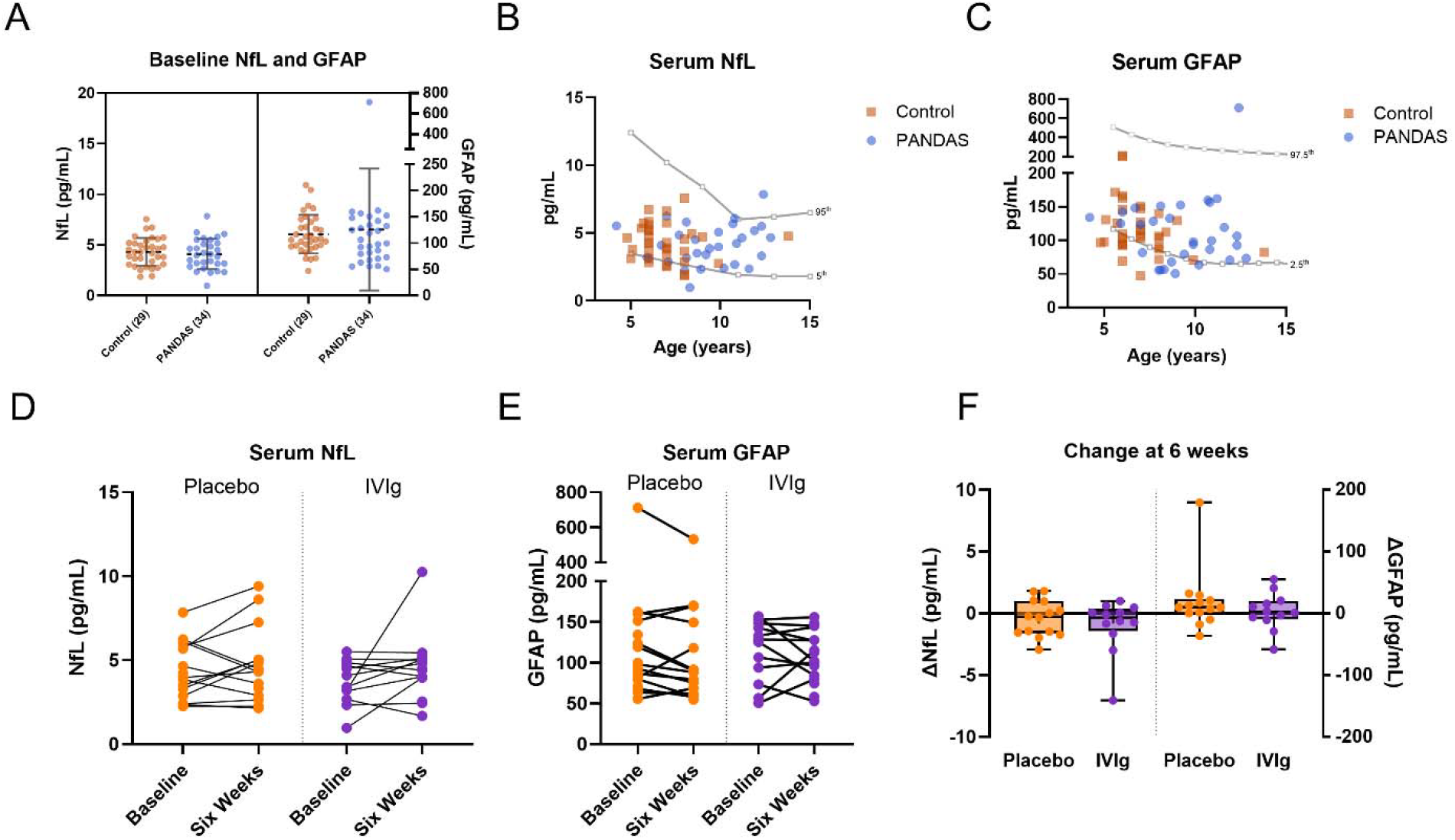
Serum neurofilament light chain (NfL) and glial fibrillary acidic protein (GFAP) in controls and participants with PANDAS at baseline and after treatment. (A) Baseline serum NfL and GFAP concentrations in controls (n=29) and PANDAS cases (n=34). Neural injury biomarker levels were comparable between groups. (B & C) Baseline serum NfL (B) and GFAP (C) plotted against age, with published pediatric age-adjusted reference percentile curves shown in gray (NfL, 5th and 95th percentiles; GFAP, 2.5th and 97.5th percentiles). (D & E) Paired serum NfL (D) and GFAP (E) measurements in PANDAS participants randomized to placebo or IVIg, obtained at baseline and 6 weeks. Neither placebo nor IVIg was associated with a consistent change in NfL or GFAP over the study interval.

Finally, we compared the NfL/GFAP ratio between groups, which has been reported to be elevated in active autoimmune encephalitis (*29*). There was no difference in the NfL/GFAP ratio between control and IVIg cases at baseline (**Supplemental Figure 8**).

### Autoantibody Screening by Whole Human Protein Microarray

We screened PANDAS CSF samples (N=25) against a ∼21,000 protein human protein array that covers ∼85% of the human proteome (HuProt v4.0, CDI Labs). For each protein spot, the median fluorescence intensity was cube-root transformed, z-scored within array, and the z-score compared to the distribution of z-scores for that protein from healthy sera control samples (N = 42). Z-scores outside of the healthy distribution were flagged as candidate autoantigens. Among PANDAS cases, there was a median of 7 candidate IgG class autoantibodies (IQR = 6, range = 1 – 38). Previously reported PANDAS autoantibodies and diagnostic autoimmune encephalitis autoantibodies were not among the candidate autoantibody set. However, one case exhibited moderate absolute reactivity to all four paralogs of the Argonaute family of proteins, AGO1, AGO2, AGO3, and AGO4 (together AGO-IgG, **Figure 4A and D**).

**Figure 4.**
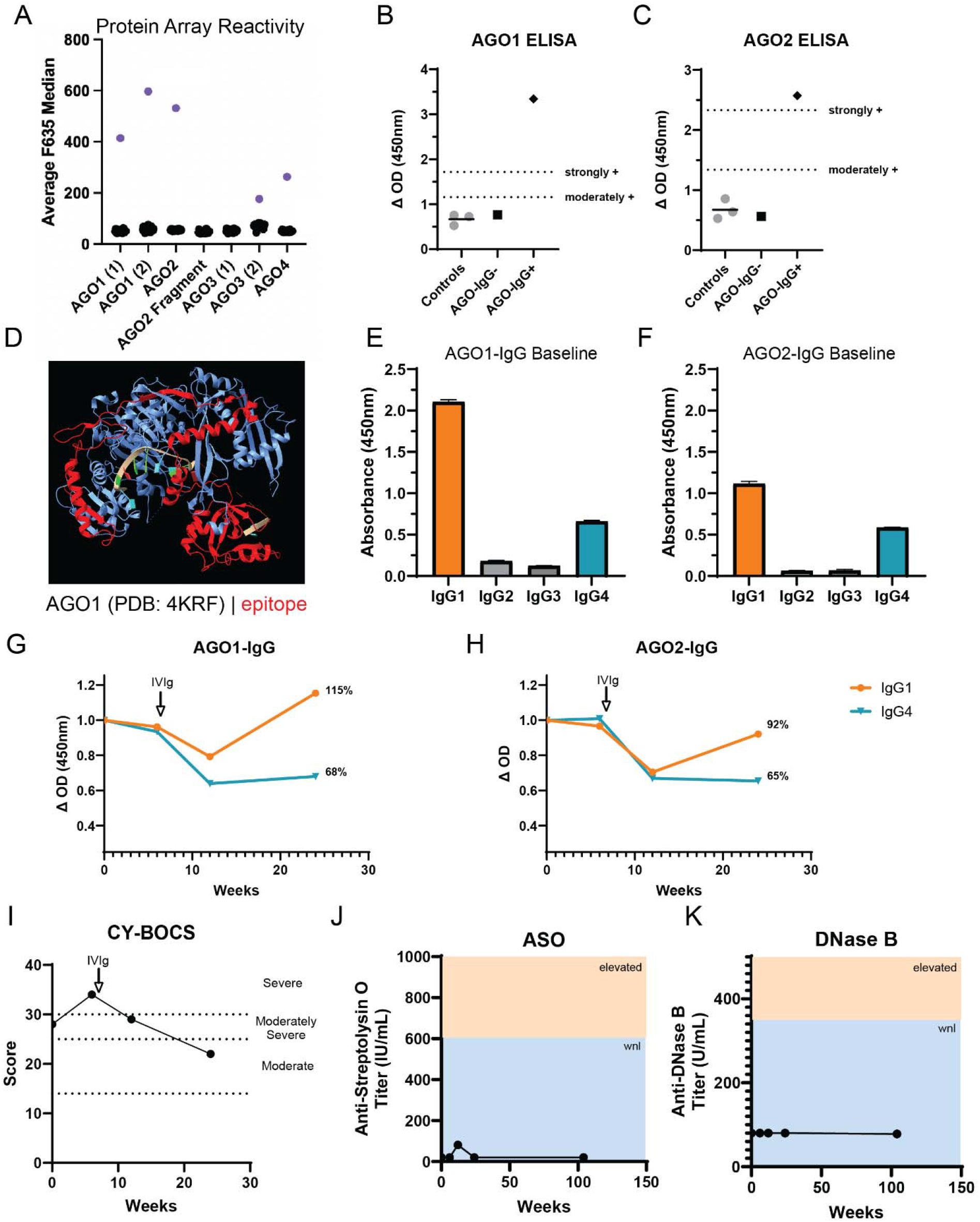
Identification and validation of Argonaute autoantibodies in a single case. (A) Screening of cerebrospinal fluid (CSF) on a HuProt human proteome microarray identified one PANDAS sample with reactivity to multiple AGO family antigens (purple points); black points represent the other PANDAS cases. (B & C) Conformation-stabilizing ELISA confirmed immunoreactivity of the index anti-AGO-positive sample (AGO-IgG+) to AGO1 (B) and AGO2 (C) relative to controls and an anti-AGO-negative comparator (AGO-IgG−). Dotted lines indicate moderate and strong positivity thresholds. (D) Region overlap among the reactive AGO proteins was used to infer a putative shared epitope, mapped in red onto the AGO1 structure (PDB: 4KRF). (E & F) Baseline subclass analysis showed that anti-AGO1 (E) and anti-AGO2 (F) reactivity was predominantly IgG1, with lower IgG4 reactivity and minimal IgG2 or IgG3 binding. (G & H) Longitudinal ELISA measurements of anti-AGO1 (G) and anti-AGO2 (H) subclass titers, shown relative to baseline, with IVIg administration indicated by the arrow. Both IgG1 and IgG4 reactivities declined after IVIg; at the last measured time point, IgG1 showed partial recovery (AGO1, 115% of baseline; AGO2, 92% of baseline), whereas IgG4 remained reduced (AGO1, 68%; AGO2, 65%). (I) Children’s Yale-Brown Obsessive Compulsive Scale (CY-BOCS) scores over the same interval, with dotted lines indicating symptom severity categories. (J, K) Antistreptolysin O (ASO) (J) and anti-DNase B (K) titers measured across the corresponding time frame and at an additional follow-up approximately 2 years later; both remained within normal limits (WNL; blue shading), with elevated ranges shown in tan.

### Validation and assessment of longitudinal AGO-IgG

AGO-IgG autoantibodies were first identified in systemic lupus erythematosus (*30*), and later in systemic autoimmune diseases more broadly (*31*). AGO-IgG was later demonstrated to be overrepresented in autoimmune sensory neuronopathy (aSNN) with or without co-morbid systemic autoimmune disease using an optimized conformation-maintaining ELISA (*32*).

We therefore recapitulated the optimized ELISA using commercial recombinant human AGO1 and AGO2 and confirmed the presence of high-titer AGO1-IgG and AGO2-IgG using previously established thresholds for conformation-stabilizing ELISA (**Figures 4B and C**) (*33*).

AGO-IgG was previously reported to be predominantly comprised of IgG1, followed by IgG4 (*34*). Therefore, we evaluated baseline serum IgG subclass reactivity to AGO1 and AGO2. In our case, we found that IgG1 was the dominant subclass against AGO1 and AGO2, followed by IgG4 (**Figures 4E and F**).

To determine the relationship between AGO-IgG and treatment, we performed a dilution series of serum samples between time points 1 (T1) – T4 and derived the change in IgG1 and IgG4 from titer estimates at each time point. For both IgG1 and IgG4, there was no change between baseline and T2 (after placebo) but a notable drop in titer at T3 (six weeks after open-label IVIg) (**Figure 4G and H**). However, whereas AGO-IgG1 rebounded at T4, AGO-IgG4 remained attenuated.

Notably, clinical labs for total IgG4 also decreased between T1 at 192.0 mg/dL (normal range, 1.0 – 108.7 mg/dL) and T4 148.0 mg/dL mg/dL (normal range, 1.0 – 121.9 mg/dL). Clinically, it was reported that the patient did not improve after 6 weeks of placebo but experienced “considerable improvement” after initiation of IVIg. Though the patient had improved 80% by parent report, the decrease in CY-BOCS was more modest. Nonetheless, symptomatic improvement corresponded to IVIg treatment (**Figure 4I**).

Finally, as measured by ELISA, AGO-IgG and symptomatic improvement had no relationship to ASO and anti-DNase B titers, which remained within the normal range during the study, including at a 2-year follow-up (**Figures 4J and K**).

## Conclusion/Discussion

Here, we evaluated the relationship between GAS-IgG and PANDAS. To do so, we used PhIP-Seq to map GAS-IgG anti-pathogen antibody repertoires in a well-curated, carefully phenotyped cohort of children diagnosed with PANDAS. We initially stratified PANDAS cases into strong, moderate, and weak GAS response groups and confirmed the suitability of PhIP-Seq by demonstrating that these groups corresponded to clinical GAS titers. Therefore, we proceeded to compare the intensity of GAS response of cases to controls but found no difference. However, in some infection-associated neurological diseases, such as multiple sclerosis (MS), the epitope specificity is more reflective of risk than seropositivity (*35*). Yet, we also found no difference in the epitope specificity of GAS-IgG between PANDAS and controls. These data indicate that a differential GAS response does not distinguish between PANDAS and unaffected children and question the utility of GAS titers for the diagnosis of PANDAS. Our data are congruent with other studies that have failed to identify a relationship between GAS-IgG and PANDAS using lower-resolution methods.

Though our data are inconsistent with GAS or anti-strep antibodies being directly causal, they do not rule out other pathogens. Therefore, we expanded our PhIP-Seq analysis to other pathogens that have been hypothesized to underlie PANS. Though we did not find a shared, PANDAS-specific pathogen response, our analysis suggested descriptive, but not significant, differences in *M*. and *C. pneumoniae* exposures in PANDAS. A limited number of studies have implicated *M. pneumoniae* in presentations with some phenotypic overlap with PANDAS, including an individual with dystonia and reversible radiographic changes in the basal ganglia (*36*), new-onset OCD in a five-year-old male that resolved after clarithromycin (*37*), and elevated *M. pneumonia* antibody titers, particularly IgA, in Tourette’s syndrome (*38*). Therefore, the potential role of *M. pneumonia* in a subset of PANDAS cases deserves further study.

We further asked whether there is evidence of neural injury in PANDAS irrespective of an identified pathogenic agent. Though at times billed clinically as autoimmune encephalomyelitis or encephalitis, T2 MRI hyperintensities, inflammatory CSF, and abnormal EEGs are uncommon in PANDAS. However, these paraclinical abnormalities are not a requirement for autoimmune encephalitis (*25*), which can further be distinguished from primary psychiatric disorders and healthy controls by serum NfL and GFAP (*29, 39, 40*). Similar to a small prior study (*41*), we found no difference in serum NfL and GFAP between PANDAS and controls. Furthermore, there was no difference in the serum NfL/GFAP ratio, which is reportedly higher in active autoimmune encephalitis as compared to MS or controls (*29*). Finally, within PANDAS, there was no difference in the change in serum NfL or GFAP levels between the placebo and IVIg groups after treatment. These data indicate that individuals who meet strict criteria for PANDAS do not have active neural injury as reflected by NfL and GFAP.

Upon antibody discovery by protein microarray, we identified AGO-IgG in one patient. Argonaute is a previously described autoantigen that is recognized as punctate cytoplasmic foci with variable nuclear staining on anti-nuclear antibody testing. The serum prevalence of anti-AGO-IgG is 0.64% in adult females (0.01% in males) (*31*), compared to 5% - 40% in inflammatory myopathies and rheumatologic disorders like systemic sclerosis (*31, 42*). More recently, AGO-IgG has been detected in the serum of patients with sensory neuropathy (6% - 17%) and CSF of those with suspected autoimmune encephalitis or paraneoplastic syndromes (6%), but was absent from the CSF of 312 healthy and noninflammatory neuropsychiatric controls (*32*). In aSNN, AGO1-IgG+ patients are more likely than AGO1-IgG-to respond to IVIg, whereas there was no differential response to steroids (*34*).

We found that our AGO-IgG+ patient had no clinical or CY-BOCS response to placebo but experienced a meaningful decrease in OCS after IVIg. We found that the patient harbored IgG1 and IgG4 class antibodies that were immunoreactive to AGO1 and AGO2. Notably, this case tested positive for anti-nuclear antibodies clinically and had elevated serum IgG4, indicative of immune dysregulation. Concordant with her symptoms, AGO-IgG titers decreased after IVIg. AGO-IgG can be suggested clinically by a positive ANA. Though this is the first definitive case of AGO-IgG in PANDAS, ANA titers above 1:200 have been reported in at least four cases of atypical OCD (*43*), and AGO-like immunostaining has been reported in at least one case of pineal germinoma-associated OCD (*44*), and two individuals with steroid-responsive OCD (*45*).

These rare cases stand in contrast to the community estimates of the prevalence of PANDAS, which approach 1 in 200. Yet the true prevalence, when following strict diagnostic criteria, is likely between 1 in 11,765 and 1 in 60,155 children (*46, 47*). Incomplete adherence to the diagnostic criteria, coupled with the high frequency of GAS infections and exposures in the pediatric population, may drive overdiagnosis of what is likely a rare clinical entity (*16*).

Overall, our data suggest that though GAS-IgG is not a unique feature of PANDAS, a subset of affected individuals may have underlying infectious exposures or autoimmunity. Importantly, syndromic PANDAS can be phenocopied by more specific genetic disorders (*48*). Syndromic PANDAS is waxing and waning, with the potential for spontaneous remission. As such, symptom-based administration of IVIg may be temporally associated with improvement despite the absence of direct benefit. Subsequent worsening of symptoms in an apparently responsive patient may lead to chronic IVIg administration or inappropriate escalation therapy, including B-cell-depleting agents. That IVIg is not universally appropriate for PANDAS is emphasized by the null results of the clinical trial from which our samples were drawn (*17*). Nonetheless, the IVIg response in one individual with AGO-IgG highlights the critical importance of biomarker-supported clinical decision making in vulnerable pediatric patients with OCS.

## Limitations

Our study was small and requires replication. Though small, our study was solely comprised of criteria-meeting PANDAS cases, whereas prior studies have shown that over half of individuals diagnosed with PANDAS in the community fail to meet criteria upon specialty reassessment (*49*). Additionally, our PhIP-Seq assay is specific to IgG, and we did not assess other anti-GAS antibody isotype responses. Nonetheless, the correlation between our PhIP-Seq-derived GAS score and clinical DNaseB and ASO titers support the validity of our PhIP-Seq measures. Additionally, PhIP-Seq assesses for antibodies that recognize linear, unmodified epitopes, where some GAS antibodies recognize conformational or post-translationally modified epitopes.

## Methods

### Participants

Patient samples in this trial were originally recruited for an IRB-approved clinical trial investigating intravenous immunoglobulin treatment for PANDAS patients (11-M-005, NCT01281969, January 2011 – August 2018) conducted at the National Institutes of Mental Health (NIMH), Bethesda, Maryland (*17*). Secondary analysis of the biospecimens and data was performed under IRB-approved protocol IRB001702. Healthy control biospecimens were enrolled under NIH IRB-approved protocol (06-M-0102, NCT00298246, February 2006 – March 2017).

### Simoa

Using SIMOA® technology from Quanterix (Billerica MA USA), Neurofilament-light chain (NfL) and Glial Fibrillary Acidic Protein (GFAP) were quantified using Neuroplex GFAP+NFL B Advantage Plus (Cat# 104670). Serum samples were diluted 1:4 in Sample Diluent supplied in the kit, clarified by centrifugation at 10,000xg, and placed into wells. Samples were then analyzed (without further onboard dilution) by acquiring duplicate results from single wells, using the SIMOA® HD-X Analyzer. Results are expressed as mean +/-Standard deviation.

### PhIP-Seq

PhIP-Seq library construction and processing of our PanSeq data have been described (*18*). Bespoke analyses for this study are described in the body of the text.

### HuProt Microarrays

Microarrays were analyzed using an in-house bioinformatic pipeline written in R, HuScore 0.7.

### Conformation-stabilizing ELISA

To validate anti-AGO immunoreactivity, we utilized a previously published ELISA protocol using recombinant human AGO1 (SinoBiological, #11225-H07B) and AGO2 (SinoBiological, #11079-H07B) proteins. Nunc MaxiSorp immunoassay plates were coated with 100 µL per well of antigen diluted to 1 µg/mL and incubated overnight at 4°C under three conditions: standard coating buffer (0.05 M carbonate/bicarbonate, pH 9.6), a conformation-stabilizing buffer consisting of 30% glycerol in coating buffer, or a denaturing condition in which antigen stock (100 µg/mL) was exposed to 0.8% SDS for 5 min and then diluted to the final coating concentration, resulting in 0.008% SDS during coating. This approach was designed to maintain denaturing conditions during the brief pre-coating step while avoiding inhibition of plate binding during antigen adsorption.

Because sera may exhibit substantial nonspecific signal, serum-specific background noise was measured for each sample using paired uncoated wells subjected to the same blocking, incubation, and wash conditions. Following blocking with 2% BSA in PBS containing 0.1% Tween-20 for 1 h at room temperature, sera were applied at 1:100 dilution in sample buffer (0.5% BSA in PBS containing 0.1% Tween-20) and incubated overnight at 4°C. Plates were then washed five times with 0.1% Tween-20 in PBS and incubated with peroxidase-conjugated donkey anti-human IgG, Fcγ specific (Jackson ImmunoResearch, #709-035-098, RRID# AB_2340494) at 1:2000 dilution, or with appropriate secondary antibodies against IgG_1_ (Thermo Fisher Scientific, Cat# A10648, RRID# AB_2534051) or IgG_4_ (Thermo Fisher Scientific, Cat# A10654, RRID# AB_2534054) subclass for 1 h at room temperature with shaking. The assay was colorimetrically developed using tetramethylbenzidine (TMB), the reaction was stopped with 0.16M sulfuric acid, and optical absorbance measured at 450 nm. Specific binding was defined as ΔOD, calculated as the optical density of antigen coated wells minus that of matched antigen uncoated wells.

For each coating condition, a positivity threshold was established as the arithmetic mean + 3 SD of healthy control ΔOD values under that same condition. To determine whether antibody binding preferentially recognized conformational epitopes, ΔOD values obtained under denaturing and stabilizing conditions were compared for each positive sample. Samples that lost ≥50% of their reactivity under denaturing relative to stabilizing conditions were classified as having conformation-dependent binding, whereas samples with <50% loss or preserved/increased reactivity under denaturing conditions were considered to recognize non-conformational epitopes.

## Acknowledgements

This research was supported by the Intramural Research Program of the National Institutes of Health (NIH). The contributions of the NIH author(s) are considered Works of the United States Government. The findings and conclusions presented in this paper are those of the author(s) and do not necessarily reflect the views of the NIH or the U.S. Department of Health and Human Services. We further acknowledge Melina V. Jones and Nicole Benoit of the NINDS Biospecimen Core Repository for their support with Simoa assays.

## Author Contributions: Conflicts of Interest

Dr. Christopher Bartley serves as a physician consultant for the non-profit Neuroimmune Foundation. Other authors declare no competing interests.

## Funding

This work was funded by NIMH ZIA awards MH002982-1, MH002982-2, and MH002982-3.

## References

1. M. Punukollu, N. Mushet, M. Linney, C. Hennessy, M. Morton, Neuropsychiatric manifestations of Sydenham’s chorea: a systematic review. Developmental Medicine & Child Neurology 58, 16–28 (2016).

2. F. R. Asbahr et al., Obsessive-compulsive and related symptoms in children and adolescents with rheumatic fever with and without chorea: a prospective 6-month study. Am J Psychiatry 155, 1122–1124 (1998).

3. A. G. Hounie et al., Obsessive-compulsive spectrum disorders in rheumatic fever with and without Sydenham’s chorea. J Clin Psychiatry 65, 994–999 (2004).

4. J. Xu et al., Antibodies From Children With PANDAS Bind Specifically to Striatal Cholinergic Interneurons and Alter Their Activity. Am J Psychiatry 178, 48–64 (2021).

5. L. Guilherme et al., Human heart-infiltrating T-cell clones from rheumatic heart disease patients recognize both streptococcal and cardiac proteins. Circulation 92, 415–420 (1995).

6. C. A. Kirvan, S. E. Swedo, L. A. Snider, M. W. Cunningham, Antibody-mediated neuronal cell signaling in behavior and movement disorders. J Neuroimmunol 179, 173–179 (2006).

7. J. L. Chain et al., Autoantibody Biomarkers for Basal Ganglia Encephalitis in Sydenham Chorea and Pediatric Autoimmune Neuropsychiatric Disorder Associated With Streptococcal Infections. Front Psychiatry 11, 564 (2020).

8. C. A. Kirvan, C. J. Cox, S. E. Swedo, M. W. Cunningham, Tubulin is a neuronal target of autoantibodies in Sydenham’s chorea. J Immunol 178, 7412–7421 (2007).

9. M. W. Cunningham, C. J. Cox, Autoimmunity against dopamine receptors in neuropsychiatric and movement disorders: a review of Sydenham chorea and beyond. Acta Physiol (Oxf) 216, 90–100 (2016).

10. H. S. Singer, C. Gause, C. Morris, P. Lopez, a. t. T. S. S. Group, Serial Immune Markers Do Not Correlate With Clinical Exacerbations in Pediatric Autoimmune Neuropsychiatric Disorders Associated With Streptococcal Infections. Pediatrics 121, 1198–1205 (2008).

11. J. F. Leckman et al., Streptococcal upper respiratory tract infections and exacerbations of tic and obsessive-compulsive symptoms: a prospective longitudinal study. J Am Acad Child Adolesc Psychiatry 50, 108–118.e103 (2011).

12. E. M. Perrin et al., Does group A beta-hemolytic streptococcal infection increase risk for behavioral and neuropsychiatric symptoms in children? Arch Pediatr Adolesc Med 158, 848–856 (2004).

13. S. E. Swedo, J. F. Leckman, N. R. Rose, From research subgroup to clinical syndrome: modifying the PANDAS criteria to describe PANS (pediatric acute-onset neuropsychiatric syndrome). Pediatr Therapeut 2, 113 (2012).

14. A. Gagliano, A. Carta, M. G. Tanca, S. Sotgiu, Pediatric Acute-Onset Neuropsychiatric Syndrome: Current Perspectives. Neuropsychiatr Dis Treat 19, 1221–1250 (2023).

15. G. Lepri et al., Clinical-Serological Characterization and Treatment Outcome of a Large Cohort of Italian Children with Pediatric Autoimmune Neuropsychiatric Disorder Associated with Streptococcal Infection and Pediatric Acute Neuropsychiatric Syndrome. J Child Adolesc Psychopharmacol 29, 608–614 (2019).

16. C. E. Helm, R. A. Blackwood, Pediatric Autoimmune Neuropsychiatric Disorder Associated with Streptococcal Infections (PANDAS): Experience at a Tertiary Referral Center. Tremor Other Hyperkinet Mov (N Y) 5, 270 (2015).

17. K. A. Williams et al., Randomized, Controlled Trial of Intravenous Immunoglobulin for Pediatric Autoimmune Neuropsychiatric Disorders Associated With Streptococcal Infections. J Am Acad Child Adolesc Psychiatry 55, 860–867.e862 (2016).

18. P. Khil, T. Esparza, C. Bartley, TIU Phage Display Platform (PanSeq) v1.0. Protocols.io, (2025).

19. G. J. Xu et al., Comprehensive serological profiling of human populations using a synthetic human virome. Science 348, aaa0698 (2015).

20. A. Olin et al., Demographic and genetic factors shape the epitope specificity of the human antibody repertoire against viruses. Nature Immunology 27, 600–612 (2026).

21. E. L. Kaplan, C. D. Rothermel, D. R. Johnson, Antistreptolysin O and anti-deoxyribonuclease B titers: normal values for children ages 2 to 12 in the United States. Pediatrics 101, 86–88 (1998).

22. C. M. Menendez et al., Dopamine receptor autoantibody signaling in infectious sequelae differentiates movement versus neuropsychiatric disorders. JCI Insight 9, (2024).

23. H. Hardin, W. Shao, J. A. Bernstein, An updated review of pediatric autoimmune neuropsychiatric disorders associated with Streptococcus/pediatric acute-onset neuropsychiatric syndrome, also known as idiopathic autoimmune encephalitis: What the allergist should know. Annals of Allergy, Asthma & Immunology 131, 567–575 (2023).

24. H. Abboud et al., Autoimmune encephalitis: proposed best practice recommendations for diagnosis and acute management. Journal of Neurology, Neurosurgery & Psychiatry 92, 757–768 (2021).

25. T. Blinder, J. Lewerenz, Cerebrospinal Fluid Findings in Patients With Autoimmune Encephalitis-A Systematic Analysis. Front Neurol 10, 804 (2019).

26. L. Tybirk, C. V. B. Hviid, C. S. Knudsen, T. Parkner, Serum GFAP - pediatric reference interval in a cohort of Danish children. Clin Chem Lab Med 61, 2041–2045 (2023).

27. S. Stukas et al., Pediatric reference intervals for serum neurofilament light and glial fibrillary acidic protein using the Canadian Laboratory Initiative on Pediatric Reference Intervals (CALIPER) cohort. Clin Chem Lab Med 62, 698–705 (2024).

28. S. Bayoumy et al., Neurofilament light protein as a biomarker for spinal muscular atrophy: a review and reference ranges. Clin Chem Lab Med 62, 1252–1265 (2024).

29. J. Piepgras et al., Glial Fibrillary Acid Protein Reflects Disease Activity in Autoimmune Encephalitis. European Journal of Neurology 32, e70207 (2025).

30. E. L. Treadwell, M. A. Alspaugh, G. C. Sharp, Characterization of a new antigen-antibody system (su) in patients with systemic lupus erythematosus. Arthritis & Rheumatism 27, 1263–1271 (1984).

31. M. Satoh et al., Characterization of the Su antigen, a macromolecular complex of 100/102 and 200-kDa proteins recognized by autoantibodies in systemic rheumatic diseases. Clin Immunol Immunopathol 73, 132–141 (1994).

32. L. D. Do et al., Argonaute Autoantibodies as Biomarkers in Autoimmune Neurologic Diseases. Neurol Neuroimmunol Neuroinflamm 8, (2021).

33. C. P. Moritz et al., Conformation-stabilizing ELISA and cell-based assays reveal patient subgroups targeting three different epitopes of AGO1 antibodies. Frontiers in Immunology Volume 13 - 2022, (2022).

34. C. P. Moritz et al., Anti-AGO1 Antibodies Identify a Subset of Autoimmune Sensory Neuronopathy. Neurology Neuroimmunology & Neuroinflammation 10, e200105 (2023).

35. C. R. Zamecnik et al., An autoantibody signature predictive for multiple sclerosis. Nat Med 30, 1300–1308 (2024).

36. J. S. Kim, I. S. Choi, M. C. Lee, Reversible parkinsonism and dystonia following probable mycoplasma pneumoniae infection. Mov Disord 10, 510–512 (1995).

37. T. E. Ercan, G. Ercan, B. Severge, M. Arpaozu, G. Karasu, Mycoplasma pneumoniae infection and obsessive-compulsive disease: a case report. J Child Neurol 23, 338–340 (2008).

38. N. Müller et al., Mycoplasma pneumoniae infection and Tourette’s syndrome. Psychiatry Res 129, 119–125 (2004).

39. M. Guasp et al., Neurofilament Light Chain Levels in Anti-NMDAR Encephalitis and Primary Psychiatric Psychosis. Neurology 98, e1489–e1498 (2022).

40. Q.-L. Lai et al., Neurofilament light chain levels in neuronal surface antibody-associated autoimmune encephalitis: a systematic review and meta-analysis. Translational Psychiatry 15, 25 (2025).

41. M. Johnson et al., No neurochemical evidence of neuronal injury or glial activation in children with Paediatric Acute-onset Neuropsychiatric Syndrome. An explorative pilot study. World J Biol Psychiatry 22, 800–804 (2021).

42. M. Ogawa-Momohara, Y. Muro, M. Satoh, M. Akiyama, Autoantibodies to Su/Argonaute 2 in Japanese patients with inflammatory myopathy. Clin Chim Acta 471, 304–307 (2017).

43. B. Pankratz et al., Cerebrospinal fluid findings in adult patients with obsessive-compulsive disorder: A retrospective analysis of 54 samples. World J Biol Psychiatry 24, 292–302 (2023).

44. W. Zhu, K. S. Burgoyne, I. M. Lesser, Presence of antibrain antibodies in obsessive-compulsive disorder secondary to pineal gland germinoma: a case report. Psychosomatics 55, 729–734 (2014).

45. D. Endres et al., Novel anti-cytoplasmic antibodies in cerebrospinal fluid and serum of patients with chronic severe mental disorders. World J Biol Psychiatry 23, 794–801 (2022).

46. E. R. Wald et al., Estimate of the incidence of PANDAS and PANS in 3 primary care populations. Front Pediatr 11, 1170379 (2023).

47. R. Goren et al., Frequency and impact of paediatric acute-onset neuropsychiatric syndrome/paediatric autoimmune neuropsychiatric disorders associated with streptococcal infections diagnosis in Canada. Paediatr Child Health 30, 157–163 (2025).

48. E. N. Dreikorn et al., Case report: Early use of whole exome sequencing unveils HNRNPU-related neurodevelopmental disorder and answers additional clinical questions through reanalysis. Frontiers in Genetics Volume 15 - 2024, (2024).

49. V. Gabbay et al., Pediatric autoimmune neuropsychiatric disorders associated with streptococcus: comparison of diagnosis and treatment in the community and at a specialty clinic. Pediatrics 122, 273–278 (2008).

