## Supplemental Methods and Figures for "Antibody Profiles in Pediatric Autoimmune Neuropsychiatric Disorders Associated with Streptococcal Infections"

**Supplemental Method for the Identification of Streptococcus pyogenes homologous peptides**

First, we searched for peptides derived from *Streptococcus pyogenes* proteins using taxonomic annotation of source proteins.

In addition, conserved peptides derived from other species may share similarity with GAS, and thus are expected to be cross-reactive. To identify such peptides, we used an unbiased BLAST search with S. pyogenes reference proteome. As a query, we downloaded the whole-genome assembly of reference strains, and extracted a set of all annotated proteins. We then used BLASTp to compare these proteins with a reference library containing peptides from all exposure library pools.

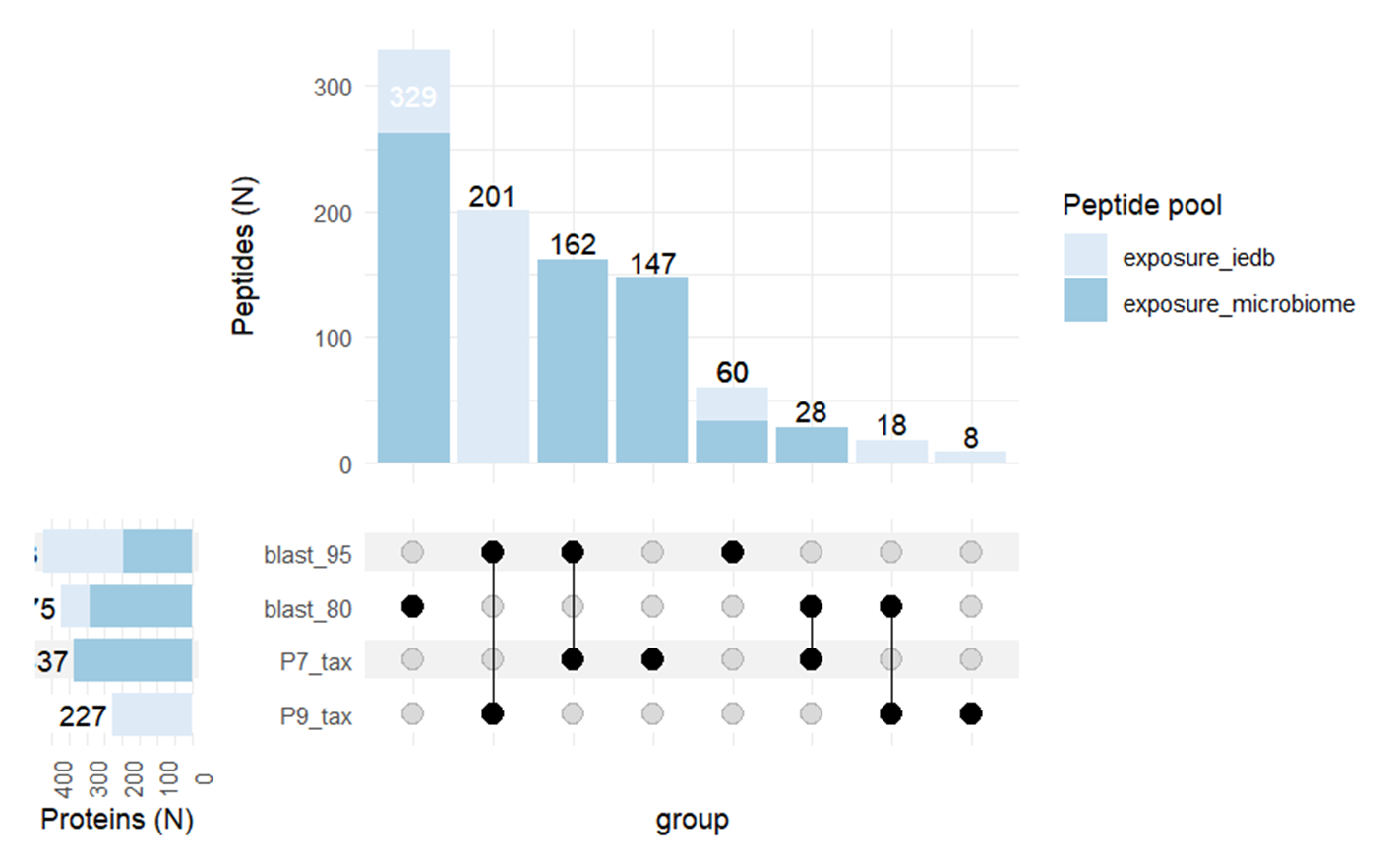

Supplementary Figure 1. Upset plot of Streptococcus pyogenes peptide representation defined in different ways, either using direct blast analysis or taxonomy retrieval.

Supplementary Figure 2. CDF plots (A) and boxplots (B) of *S .pyogenes* aggregated peptide reactivity score calculated for study samples and set of matching mock IPs. GAS reactivity score was calculated by summing over all GAS peptides log2-transformed normalized reactivity values. There is strong separation of GAS scores for patient samples and mock IPs.

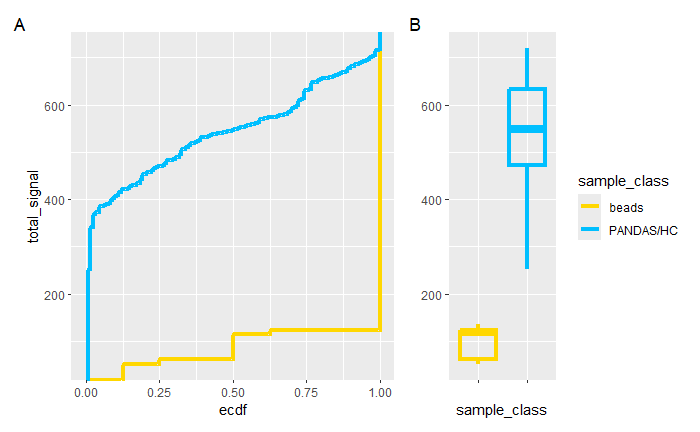

Supplementary Figure 3. Correlation between GAS score and clinical test titers.

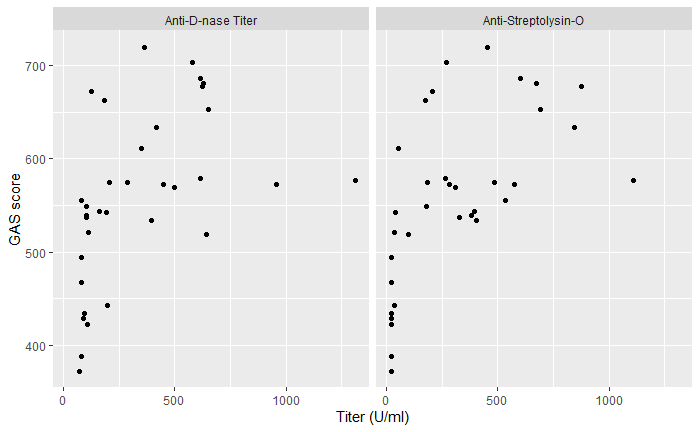

Supplementary Figure 4. CDF plots (A) and boxplots (B) of MP aggregated peptide reactivity score calculated for study samples and set of matching mock IPs. MP reactivity score was calculated by summing over log2-transformed normalized reactivity values. (C) Heatmap of re-centered peptide reactivities (max_value >2).
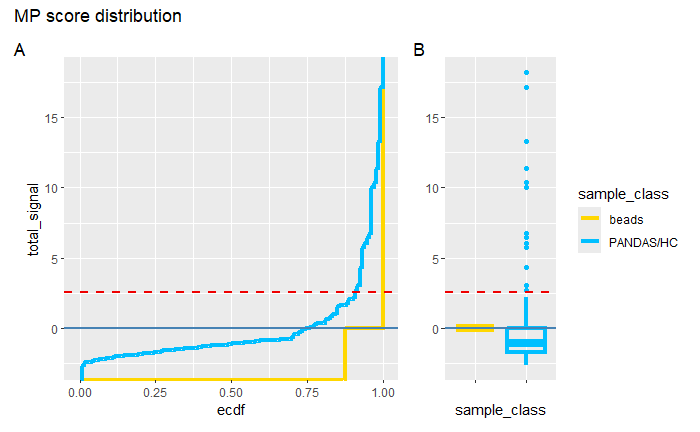

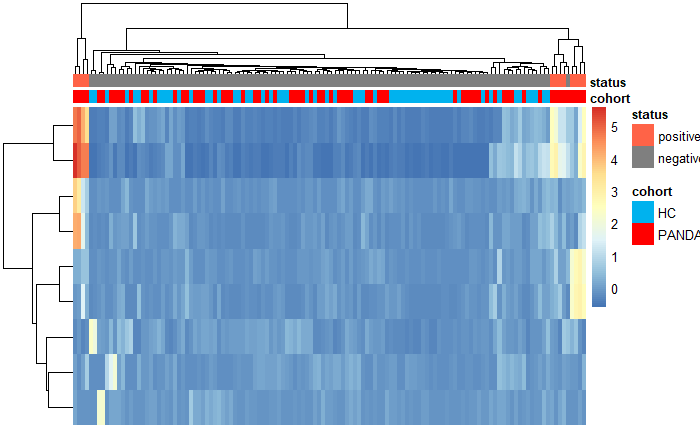

Supplementary Figure 5. CDF plots (A) and boxplots (B) of CP aggregated peptide reactivity score calculated for study samples and set of matching mock IPs. CP reactivity score was calculated by summing over reactive log2-transformed normalized reactivity values. (C) Heatmap of re-centered peptide reactivities (max_value >2).
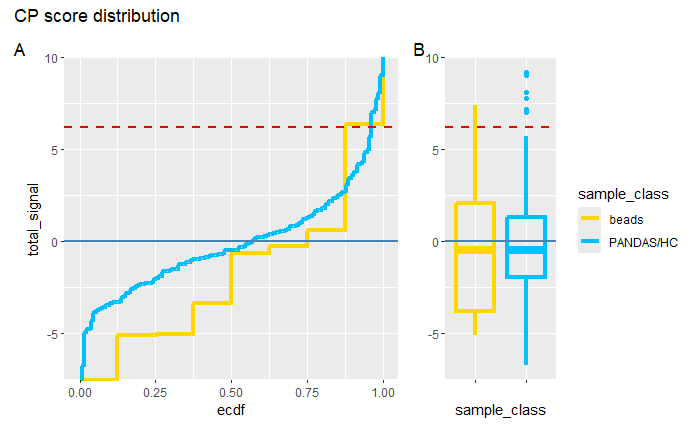

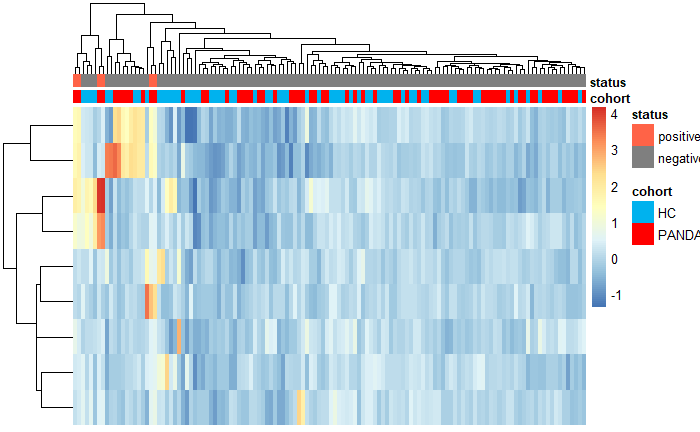

Supplementary Figure 6. CDF plots (A) and boxplots (B) of HSV aggregated peptide reactivity score calculated for study samples and set of matching mock IPs. HSV reactivity score was calculated by summing over log2-transformed normalized reactivity values. (C) Heatmap of re-centered peptide reactivities (max_value >2).
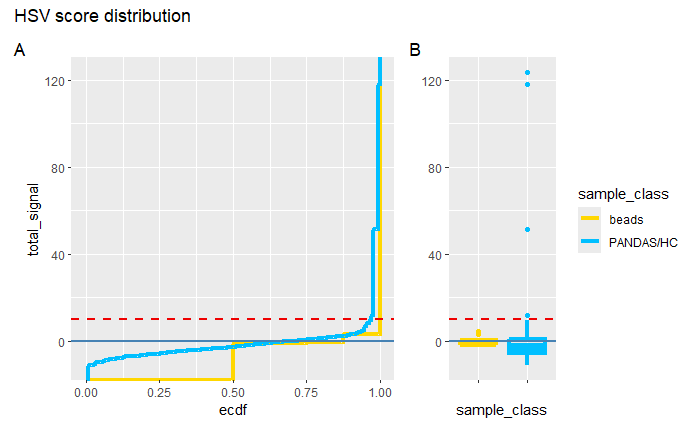

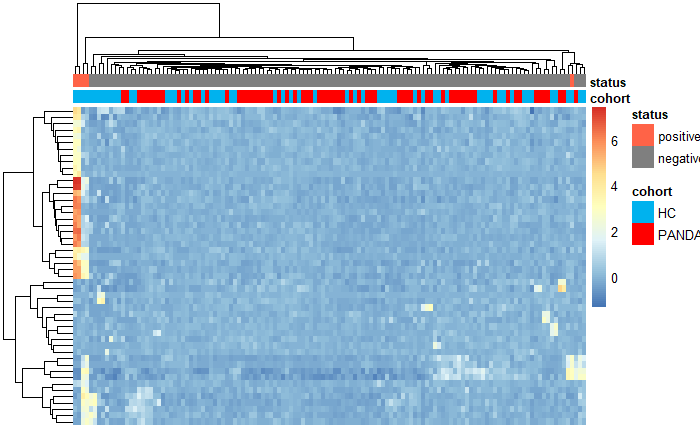

Supplementary Figure 7. CDF plots (A) and boxplots (B) of EBV aggregated peptide reactivity score calculated for study samples and set of matching mock IPs. EBV reactivity score was calculated by summing over log2-transformed normalized reactivity values. (C) Heatmap of re-centered peptide reactivities (max_value >2).
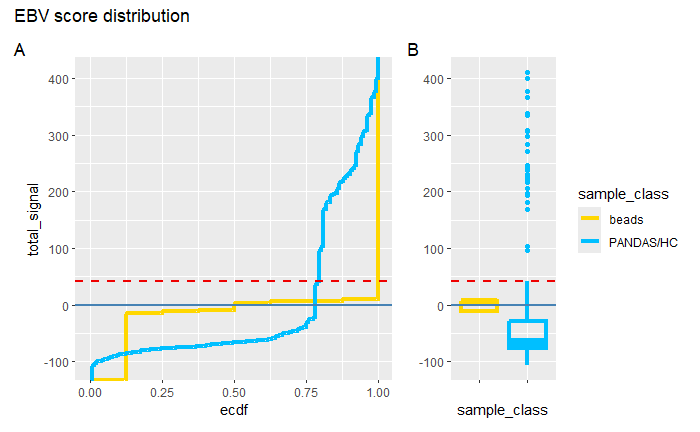

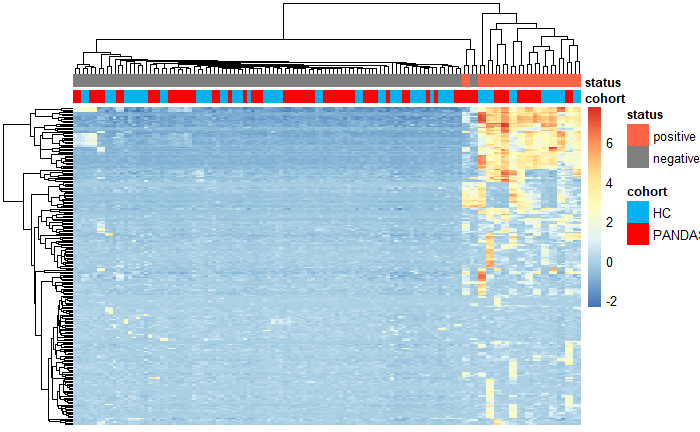

Supplementary Figure C. CDF plots of age distribution for PANDAS and HC comparator cohort. (A) Full range of ages. (B) Only patients younger than 10 included.

A

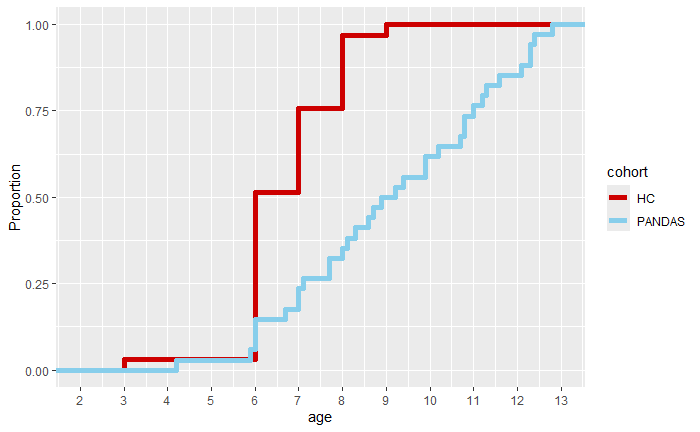

B

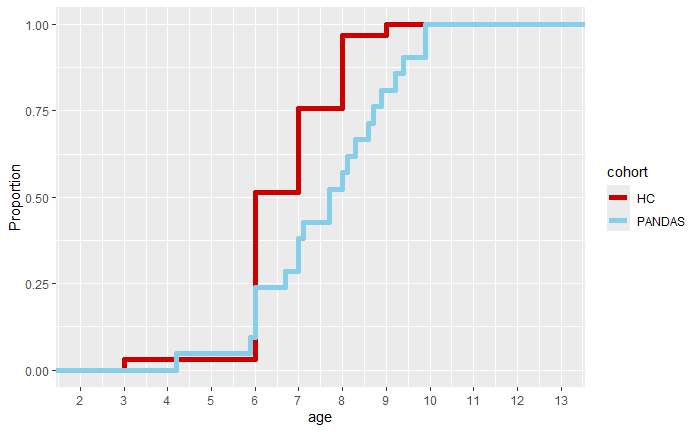

Supplemental Figure 8. Baseline NfL/GFAP ratio in sera from controls and PANDAS cases

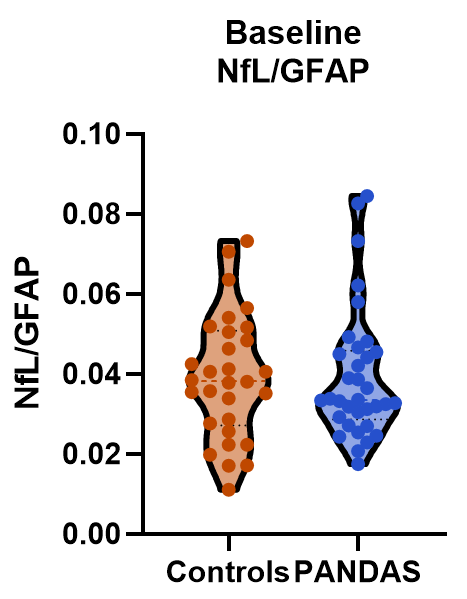
